# Inferring identical by descent sharing of sample ancestors promotes high resolution relative detection

**DOI:** 10.1101/243048

**Authors:** Monica D. Ramstetter, Sushila A. Shenoy, Thomas D. Dyer, Donna M. Lehman, Joanne E. Curran, Ravindranath Duggirala, John Blangero, Jason G. Mezey, Amy L. Williams

**Affiliations:** Department of Biological Statistics and Computational Biology, Cornell University, Ithaca, NY 14853, USA; South Texas Diabetes and Obesity Institute, University of Texas Rio Grande Valley, Brownsville, TX 78520, USA and Edinburg, TX 78539, USA; Department of Medicine, University of Texas Health San Antonio, San Antonio, Texas 78229, USA; Department of Genetic Medicine, Weill Cornell Medicine, New York, NY 10065, USA

## Abstract

As genetic datasets increase in size, the fraction of samples with one or more close relatives grows rapidly, resulting in sets of mutually related individuals. We present DRUID—Deep Relatedness Utilizing Identity by Descent—a method that works by inferring the identical by descent (IBD) sharing profile of an ungenotyped ancestor of a set of close relatives. Using this IBD profile, DRUID infers relatedness between unobserved ancestors and more distant relatives, thereby combining information from multiple samples to remove one or more generations between the deep relationships to be identified. DRUID constructs sets of close relatives by detecting full siblings and also uses a novel approach to identify the aunts/uncles of two or more siblings, recovering 92.2% of real aunts/uncles with zero false positives. In real and simulated data, DRUID correctly infers up to 10.5% more relatives than PADRE when using data from two sets of distantly related siblings, and 10.7–31.3% more relatives given two sets of siblings and their aunts/uncles. DRUID frequently infers relationships either correctly or within one degree of the truth, with PADRE classifying 43.3–58.3% of tenth degree relatives in this way compared to 79.6–96.7% using DRUID.

## Introduction

Pedigree relationships are fundamental to genetics, with segments of each individual’s genome necessarily transmitted across successive generations of ancestors in order to reach the present day. The dynamics of inheritance within pedigrees not only underlie each person’s genetic heritage, they affect observed variation in ways that are not fully captured by most population genetic models^1^. The inappropriateness of traditional models has generally had limited impact on analyses to date, largely due to the low rates of closely related samples in most prior studies. Recently however, numerous efforts have been undertaken and are ongoing to generate genetic datasets on a massive scale^2–5^. These substantial studies provide opportunities to better understand human disease genetics^3–6^ and to characterize fine-scale population structure and recent demography^7^. Yet to perform these and effectively all genetic analyses, it is necessary to account for the relatedness structure within the sample data^1,8–10^.

Given these considerations, relatedness estimation is becoming both more important and more precise because of the widespread relatedness in large samples. A striking illustration of the scope of this is in the ~500,000 sample UK Biobank data in which almost one-third of the individuals have a third degree (e.g., first cousin) or closer relative in the cohort^2^. While this is a higher rate of close relatives than expected by random sampling^2^, the proportion of individuals with one or more close relatives in a dataset grows rapidly with its size, eventually such that all samples have these relationships.

While association studies account for relatedness using kinship estimates that do not require knowledge of pedigree relationships^11,12^, population genetic analyses have the potential to benefit from explicit modeling of segregation patterns using known meiotic distances. As an example, family-based phasing and imputation can achieve near perfect accuracy, with trio phasing used as the gold standard to evaluate accuracy in population phasing methods^13^. Besides these inferences, pedigree samples and their relationships are needed to perform studies of *de novo* mutation^14–16^ and recombination^17–20^. Thus, recovering pedigree relationships from genetic data has the potential to empower population genetic analyses in general, and to enhance studies of the two fundamental sources of genetic variation through mining existing datasets for the required family data.

In order to make use of the multi-way relatedness present in large samples, a key question is how best to combine information among the individuals. Two recently developed methods use composite likelihood approaches that enable them to outperform pairwise relatedness inference^21,22^. Results from these methods demonstrate that leveraging multi-way relatedness signals does improve accuracy, but their reliance upon composite likelihoods may be suboptimal. In order to learn the best strategies for relatedness inference, we recently analyzed the accuracy of 12 pairwise relatedness methods and found that: (1) identical by descent (IBD) segment-based methods perform best for classifying a broad range of degrees of relatedness, and (2) the overall accuracy of all methods is highest for close relationships^23^.

Building on these findings, we present DRUID, Deep Relatedness Utilizing Identity by Descent, a method that infers relatedness between distant relatives and an ungenotyped ancestor of a set of close relatives. DRUID leverages the fact that, besides the descendants of an individual, all relatives arise through a relationship with at least one parent of each sample. Indeed, a given individual’s parents necessarily transmit all the genetic segments they co-inherited with their non-descendant relatives—the so-called IBD segments between these individuals^24^. Yet because a parent transmits only half of his or her genome to each child, and because this transmission is random, the amount of DNA shared with a distant relative has non-trivial variance, with an increasing coefficient of variation for more distant relatives^25^. In consequence, and consistent with results from real data^23^, relatedness inference accuracy improves by considering the IBD sharing of a parent with some distant relative.

DRUID leverages siblings and uses a new approach to identify aunts and uncles of a set of siblings, enabling inference of the IBD sharing of an ungenotyped grandparent. The method for locating aunts and uncles works on the basis that such individuals, being full siblings of their nieces/nephews’ parent, necessarily share a sizable amount of DNA that is IBD on both haplotype copies with that (ungenotyped) parent. Using data from two siblings, this method recovers 92.2% of real aunts and uncles (90.1% in simulations) with no false positives.

When sampling of relatives is very dense, there may be two sets of close relatives that are mutually related to one another. In this case, DRUID performs inference of the IBD sharing between two ungenotyped individuals that are the respective ancestors of the two close relative sets. This removes up to four generations of distance between the individuals whose relationships are to be inferred, yielding substantial accuracy gains.

We compared DRUID to the multi-way relatedness method PADRE^21^ and to a pairwise inference approach that uses IBD segments called by Refined IBD^23,26^. Both PADRE and DRUID perform much better than pairwise analysis: using simulated data for two sets of full siblings that are distantly related to each other, the multi-way methods classify 5.0–35.4% more samples to their exact degree of relatedness. DRUID is also more accurate than PADRE when using siblings or siblings and aunts/uncles, with up to 9.2–29.6% more real and simulated relatives inferred exactly, and with greater accuracy in nearly all these analyses. Turning to relatedness inference that is within one degree of the truth—a metric that is most relevant for more distant relationships—DRUID detects 79.6–96.7% of simulated tenth degree relatives while PADRE classifies 43.3–58.3% of such relatives in this way. DRUID utilizes siblings (and detected aunts/uncles) for most of its improvement whereas PADRE attempts to utilize a wider range of relationships. Accordingly, we performed analyses that included either no siblings or one pair of distantly related siblings. In 14 out of 15 of these scenarios, results were mixed, with PADRE inferring up to 6.7% more fourth through sixth degree pairs, and DRUID classifying up to 18.3% more of these relatives correctly. PADRE’s accuracy is reduced for seventh through tenth degree relationships in these scenarios, with DRUID correctly inferring 5.8–38% more such relatives. DRUID is freely available and open source.

## Methods

DRUID begins by taking input IBD segments inferred from genotype data and performs relatedness inference in two stages. First, it infers the pedigree structure of samples that are connected through first degree relationships—relationships that are very likely to be inferred correctly^23^. In cases where DRUID identifies two or more siblings and only one or neither of their parents, it also locates aunts and/or uncles of these siblings. Second, DRUID performs its primary relatedness inference algorithm, combining IBD information from individuals in the inferred pedigrees when possible to calculate the expected genome-wide IBD sharing proportion between an ungenotyped ancestor and a more distant relative (Figure 1A). Using this quantity, the method then infers the likely degree of relatedness between that ancestor and the distant relative—who may also be an ungenotyped individual. When the relationship to the distant relative arises through one of the close relatives’ ancestors (rather than through descent from one of these individuals), the ungenotyped ancestor will be more closely related to the distant sample than the genotyped individuals are.

**Figure 1:**
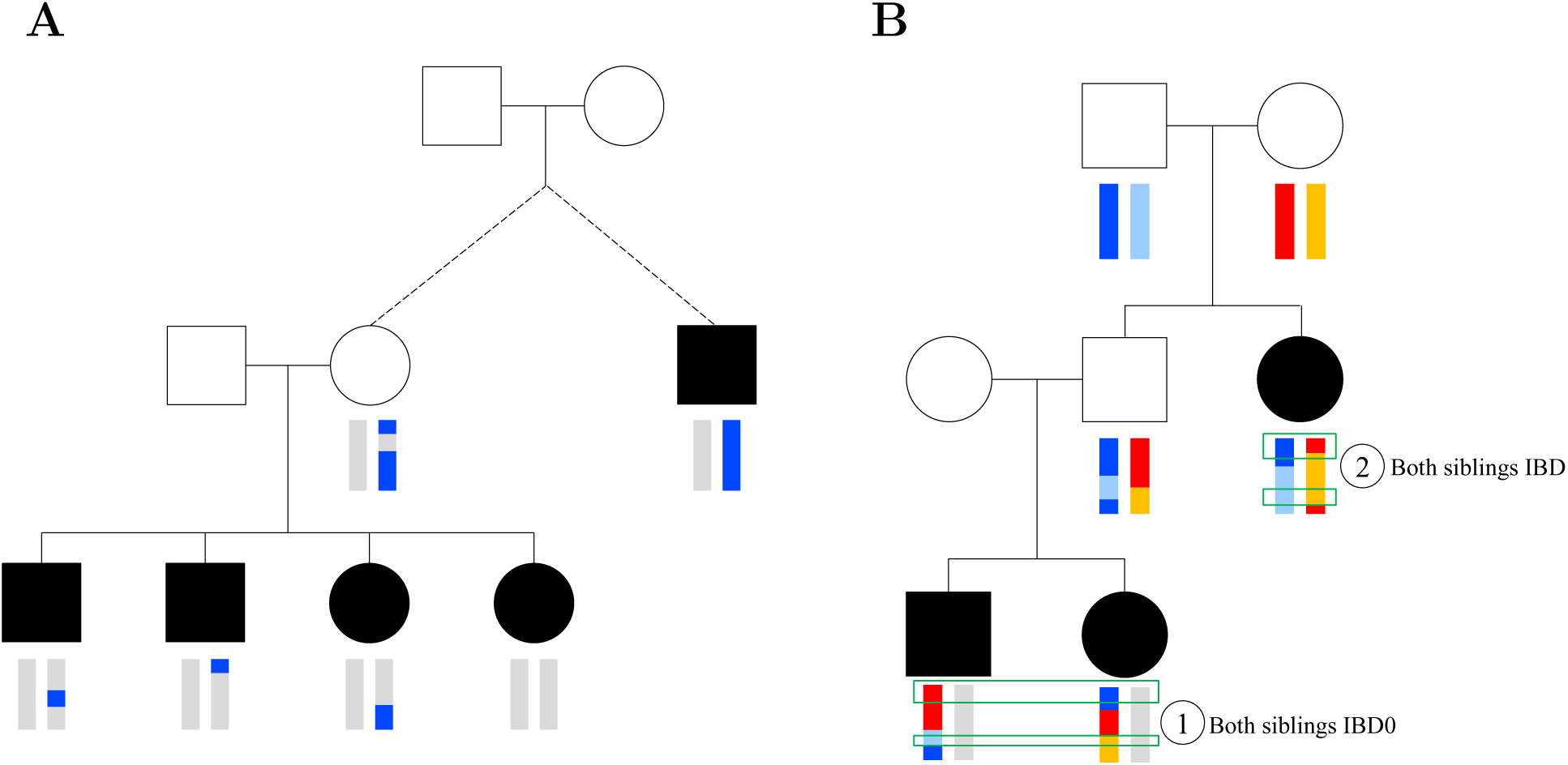
(**A**) Pictorial depiction of DRUID’s relatedness inference approach. Genotyped individuals are shown as filled shapes and haplotypes are colored vertical bars below analyzed samples; the dashed line indicates the number of generations to the most recent common ancestors between the full siblings and the distant relative on the right is unknown. The blue regions in the full siblings represent IBD segments shared with the distant relative on the right. DRUID infers the ungenotyped mother’s IBD profile as the union of her children’s IBD segments. (**B**) Example haplotype transmissions from grandparents to two full sibling grandchildren (bottom generation) as well as the siblings’ father and aunt, with regions of the same color descended from the corresponding grandparent chromosome (we ignore the gray chromosomes for simplicity). Given genotype data for the two siblings and their aunt, the IBD^(011)^ regions are those where the two siblings are (1) IBD0 with each other, and (2) both are IBD with the aunt, indicated by green boxes. These are positions where the siblings’ ungenotyped father is IBD2 with the aunt.

Because most genotype datasets contain samples that were only collected relatively recently, we make the assumption that the distant relatives do arise through an ancestor and not via descent. DRUID also assumes that there are no errors in the detected IBD segments and that there is no consanguinity among either the close or more distant relatives. That is, we assume that the two parents of any set of siblings are not related to each other and that all IBD segments shared with a sample not in a set of close relatives descend from only one of the close relatives’ common ancestors. We have found that, although errors in detected IBD segments do occur, their overall effect is limited when using a relatively accurate IBD detection method. We implemented DRUID to utilize IBD segments detected using Refined IBD^26^, but the approach is generally applicable to any method that reports whether samples share one or two IBD segments at a given position. Throughout, we refer to regions in which two individuals share zero, one, or two IBD segments with each other as IBD0, IBD1, and IBD2, respectively. We also use the term *distant relative* to refer to a sample that is not a member of a given set of close relatives and has third degree or more distant relatedness to at least one individual in that set. For first and second degree relatives, DRUID performs inference using the inferred pairwise kinship coeffcient 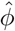 (below). DRUID infers up to 13th degree relatedness between any pair it considers, including ungenotyped individuals, labeling others as unrelated. (Most 13th degree relatives share no IBD regions; however, our simulations suggest that ~10% do share at least one ≥ 2 cM segment, so we attempt this inference.) DRUID propagates relatedness degrees to descendant individuals in the pedigree structures it infers, including more distant than 13th degree relatedness for these descendants.

### Inferring pedigree structures

To infer the sets of closely related samples and their pedigree structures, DRUID generates a graph in which nodes represent samples and edge labels indicate the relationship type between the linked pair. The input IBD segments are informative about the relationships between the samples, and we use these to estimate 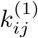 and 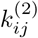, the proportion of their genome that samples *i* and *j* share IBD1 and IBD2, respectively. These estimates are simply the sum of the lengths of the IBD1 and IBD2 segments shared between *i* and *j* divided by the total genome length, with all lengths in genetic units (e.g., centiMorgans [cM]). From this, we derive the estimated kinship coefficient between *i* and *j* as 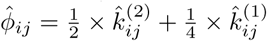 and deduce their likely relationship type based on this coeffcient and 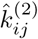 using the values in Table 1. DRUID uses this pairwise kinship estimate to infer first and second degree relatives, to reconstruct pedigrees, and to infer relatedness between samples not in pedigrees. Initially in pedigree reconstruction, the method considers only parent-child, full sibling, and monozygotic (MZ) twin relationships. In the case of MZ twins, DRUID analyzes IBD information from only one of the twins and provides relatedness estimates for the omitted twin(s) that are identical to the analyzed individual.

**Table 1:** Relationship classification rules used by DRUID and DRUID*_C_*. The ranges of 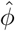 and their mappingto relationships are based on those used by KING^27^. First degree relatives that do not fall into a given 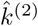 range are not used in DRUID’s pedigree reconstruction. MZ twin: monozygotic twin. * DRUID uses a lower 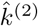 value (and consequently 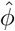) for full siblings than that suggested in the KING paper as this yields better results in simulations.

|  | DRUID |  | DRUID<sub>C</sub> |  |
| --- | --- | --- | --- | --- |
| | $\hat{k}^{(2)}$ | $\hat{\phi}$ | $\hat{k}^{(2)}$ | $\hat{\phi}$ |
| MZ twin | $[\frac{1}{2^{1/2}}, 1]$ | $[\frac{1}{2^{3/2}}, 1)$ | $[\frac{1}{2^{1/2}}, 1]$ | $[\frac{1}{2^{3/2}}, 1)$ |
| Full sibling* | $[\frac{1}{2^{11/4}}, \frac{1}{2^{3/2}})$ | $[\frac{1}{2^{7/2}}, \frac{1}{2^{3/2}})$ | $[\frac{1}{2^{5/2}}, \frac{1}{2^{3/2}})$ | $[\frac{1}{2^{5/2}}, \frac{1}{2^{3/2}})$ |
| Parent-child | $[0, \frac{1}{2^{9/2}})$ | $[\frac{1}{2^{5/2}}, \frac{1}{2^{3/2}})$ | $[0, \frac{1}{2^{9/2}})$ | $[\frac{1}{2^{5/2}}, \frac{1}{2^{3/2}})$ |
| Double cousin | $[\frac{1}{2^{9/2}}, 1]$ | $[\frac{1}{2^{7/2}}, \frac{1}{2^{5/2}})$ | $[\frac{1}{2^{9/2}}, 1]$ | $[\frac{1}{2^{7/2}}, \frac{1}{2^{5/2}})$ |
| $D^{\text{th}}$ degree | - | $[\frac{1}{2^{(2D+3)/2}}, \frac{1}{2^{(2D+1)/2}})$ | - | $[\frac{1}{2^{(2D+3)/2}}, \frac{1}{2^{(2D+1)/2}})$ |

Starting with an empty graph, DRUID adds nodes and edges corresponding to all inferred full sibling relationships. Following this, the method ensures that for all connected components, the nodes contained in it are all directly connected to one another as full siblings. If any member does not have a full sibling relationship with another individual in the component, DRUID checks whether the majority (rounded up in cases of even numbers) of pairs are labeled as full siblings of that individual. In that case, it labels any unconnected pairs as full siblings as well, and otherwise, it breaks the minority full sibling connections. DRUID can also be run so that close relative inferences are made more conservatively, which we refer to as DRUID*_C_*. In this mode, for non-fully connected components, DRUID successively removes edges to nodes that are the least connected (randomly selecting a node in the case of a tie) until the resulting components are fully connected. While we expect the standard DRUID method to be applicable in most settings, DRUID*_C_* may be useful in datasets where less common relationship types exist, such as three-quarter siblings.

Next, DRUID incorporates parent-child relationships into the graph, and, when possible, determines which individual is the parent using analysis of relatedness to full siblings. Full sibling relationships are informative about which sample is the parent because full siblings have the same parents. Suppose *i* and *j* are inferred to have a parent-child relationship. If *j* has full siblings in the graph, and if one has a parent-child relationship to *i*, DRUID assigns *i* as the parent of all full siblings of *j*. Alternatively, if none of the full siblings of *j* have a parent-child relationship with *i*, DRUID assigns *i* as the child of *j*. When neither *i* nor *j* have full siblings, DRUID labels *i* and *j* as a parent-child pair without assigning either as the parent.

As described below, DRUID is able to infer aunts and uncles of a set of siblings, but because mistakes in the inferred pedigree structures can lead to systematic errors in later classification, we do not currently infer half-siblings or other types of relatives. However, DRUID does utilize user-specified relationships, including half-siblings, grandparent-grandchild pairs, and known directions for parent-child relationships. To incorporate this information, DRUID performs initial inference of a pedigree using first degree relationships (above) and then connects second degree relatives with non-specific second degree relationship edge types. It then compares the inferred pedigree with the user specifications, prints a warning for any discrepancies, and changes the relationships to match the specifications. Following this, the algorithm proceeds to infer aunts and uncles.

### Identifying aunts and uncles of a set of siblings

In principle, the length of IBD segments shared between second degree relatives are informative about their underlying relationship type: grandparent-grandchild, half-sibling, avuncular, or double cousin. However, the method RELPAIR, which implements an approach based on this idea, has limited ability to discriminate between these relationships, with previously reported classification true positive rates ranging from 37% to 72% for the different relationship types (excluding double cousins which it does not consider)^28^.

In DRUID, we take a different approach based on the IBD sharing patterns among a set of three samples consisting of a pair of full or half-siblings and a second degree relative, determining whether that relative is an aunt or uncle of the siblings. A distinguishing feature of full siblings is that they share a substantial fraction of their genome IBD2 with each other. As such, the ungenotyped parent of a pair of siblings will share some regions IBD2 with the siblings’ aunt or uncle but not with other types of second degree relatives. This marks a unique sharing pattern that allows us to pinpoint these relatives with high specificity. As depicted in Figure 1B, at positions in which two siblings share no IBD segments with each other or are IBD0, they will have inherited distinct haplotype copies from their parents. If the two siblings each also share an IBD segment with another relative in these regions, one of their parents must share two distinct haplotypes IBD with that relative. (This ignores double cousins to whom both parents are related, a case we address below.) An appreciable level of this IBD pattern among the three samples is a strong indicator that the second degree relative is a full sibling of an ungenotyped parent and therefore an aunt or uncle of the two siblings.

To infer aunts and uncles, DRUID locates all pedigrees containing two or more full or half-siblings that lack data for one or both parents. It then finds all samples that are inferred as second degree relatives of at least one of these siblings (Table 1) and considers the 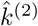 level between each sibling and that relative. Double cousins are second degree relatives that are expected to share 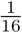 of their genome IBD2 with each other, and these relatives can also have non-trivial proportions of the IBD pattern otherwise indicative of aunts and uncles. DRUID infers any relative *r* with whom one of the siblings *s* shares 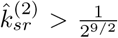 as a likely double cousin, and it ignores these in later analyses (Figure 2A, S1A).

**Figure 2:**
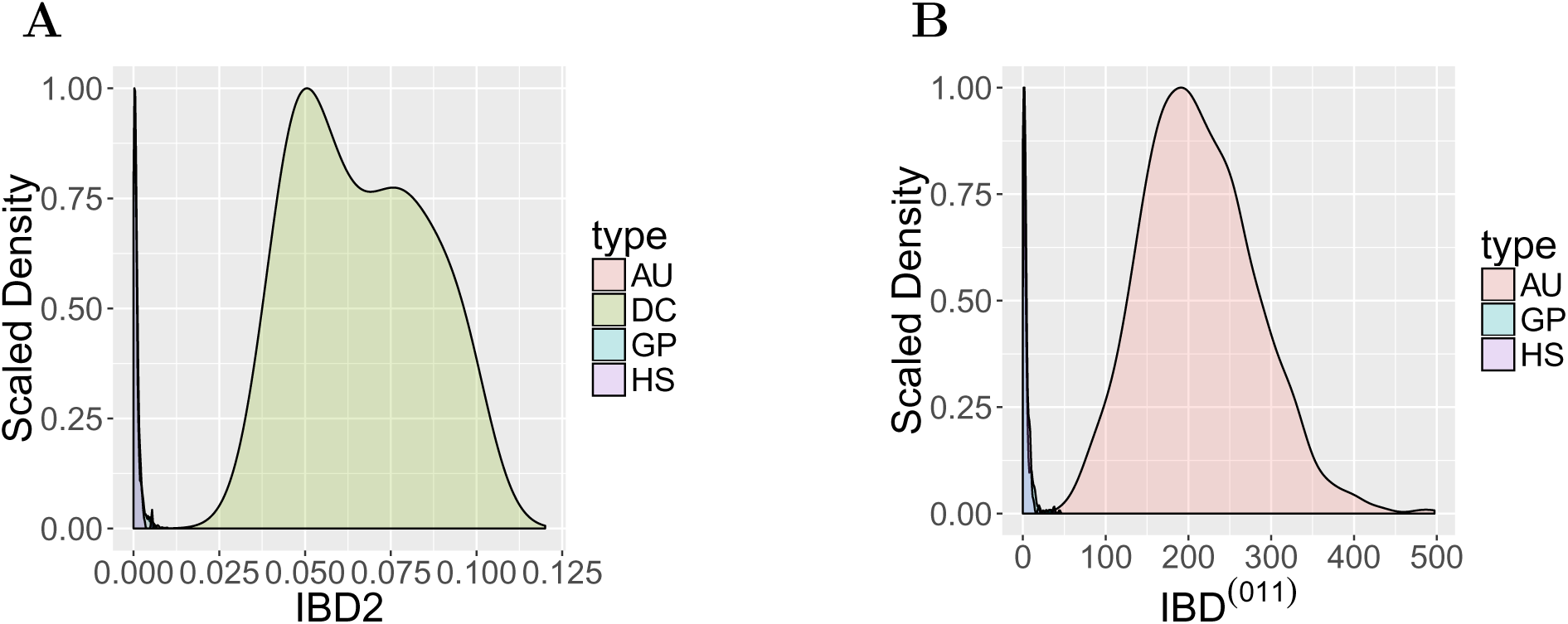
(**A**) Scaled density showing the genome proportion shared IBD2 between real second degree relative pairs from the SAMAFS data. Abbreviations: AU, aunt/uncle of a given sample; DC, double cousins; GP, grandparent of a given sample; and HS, half-siblings. (**B**) Length of genome found to be IBD^(011)^ between two full siblings and various types of second degree relatives using real data from SAMAFS. Abbreviations for the relationship between the siblings and the indicated relative as in (A). Double cousins are filtered based on their IBD2 proportion and therefore not shown. Plot colors are translucent with most relationship types overlapping one another near 0 in both panels.

For the remaining second degree relatives, the method first randomly selects a pair of siblings *s*_1_ and *s*_2_ and calculates the total genetic length of the regions they are inferred to be IBD0 with each other and where both are inferred to be IBD with the other relative *r*. (We count both IBD1 and IBD2 sharing between each sibling and *r*, though the total IBD2 levels tend to be small.) This gives a value we refer to as 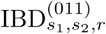, which is a lower bound on the length of the IBD2 sharing between *r* and the parent of *s*_1_ and *s*_2_ (assuming all the IBD segments descend from only one parent). Analyses of both real and simulated second degree relatives indicate that when 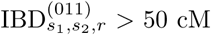, the second degree relative is effectively always an aunt or uncle of the siblings (Figure 2B, S1B). These data also indicate that when 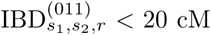, the second degree relative is very unlikely to be an aunt/uncle. The algorithm makes inference using these length values as cutoffs, and when 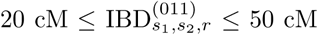 for a given sibling pair, it moves to consider a different pair of siblings to get more definitive results. If analyses of all sibling pairs fall in this ambiguous range, the relative is not inferred as an aunt/uncle. Upon inferring an individual as an aunt/uncle of a set of siblings, DRUID adds her/him to the graph as the indicated relationship, and also incorporates avuncular connections with any known full siblings of that aunt/uncle. If the input specifications include avuncular relationships, DRUID compares these to the inferred avuncular relationships at this stage, and it prints a warning if it did not infer any input relationship. In all cases, it trusts the user specifications and performs later inference using these.

With the pedigree relationships between sets of close relatives determined, DRUID undertakes the second stage of its analysis by reconstructing the IBD profiles of the ancestors of these sets. This inference focuses on two configurations of close relatives: siblings (including user-specified half-siblings) and siblings together with their aunts/uncles. As discussed later, DRUID also uses a heuristic to distinguish parent-child pairs when pedigree reconstruction fails to identify which individual is the parent.

### Inferring IBD sharing of a parent using data from siblings

A parent transmits to each child a random portion of the IBD regions he/she shares with any relative. Whereas a single child inherits only half of each parents’ genome, data for additional children include a more complete subset of the genomes of their parents, including receiving a larger fraction of the IBD regions they each carried. In particular, If 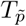 corresponds to the amount of DNA a parent 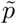 transmitted to a set *S* of full sibling children, then assuming the haplotype transmitted at each position follows a binomial distribution with Mendelian (50%) transmission, 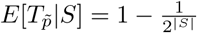.

Given the assumption that the parents of *S* are unrelated, only one parent will have transmitted all the IBD segments that those siblings share with any distant relative *d*. Thus, although genetic data for the two parents are unobserved, the union of all IBD segments shared by the siblings with *d* constitutes a partial set of the IBD regions one of the parents shared with *d* (Figure 1A, S2A). Notably, which parent transmitted these IBD segments is unknown, but this information is not needed to determine the degree of relatedness between that ungenotyped parent 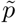 and *d*.

We relate the proportion of the genome that an ungenotyped parent 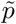 and *d* share IBD1 to the observed IBD segments in *S* as:

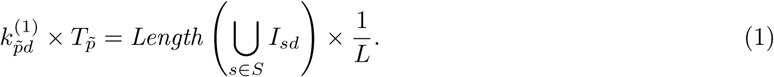

Here, *I_sd_* contains the regions that are detected as IBD between a given sibling *s* and *d*, where we include both IBD1 and IBD2 segments—the latter arising infrequently and due to error when our assumptions hold. The *Length*(*I*) function gives the genetic length of the IBD regions in *I*, and *L* is the total length of the genome, both in cM. As the union of all IBD regions in the siblings contain only a proportion 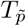 of the parent’s IBD regions, we scale the parent’s IBD sharing proportion 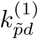 by this quantity. This implicitly models the sharing proportion in the unobserved regions of the parent’s genome as being the same as in the transmitted regions.

Because our aim is to provide an estimate of the degree of relatedness between all samples, we compute a point estimate of 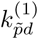 by calculating its expectation rather than modeling its full distribution. Conveniently, 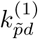 (the fraction of DNA 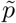 shares with *d*) is independent of 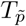, so dividing the right hand side of Equation 1 by 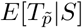 provides this expectation. The estimated kinship coeffcient between 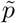 and *d* is then 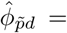 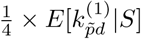, from which we infer a degree of relatedness (Table 1).

An alternative to dividing by the expectation of 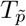 is to estimate its realized value by analyzing the observed IBD sharing between the siblings. Specifically, at positions where full siblings are all IBD2 with one another, both parents will have transmitted only one haplotype copy, or half of their genome. Conversely, at positions where at least two children are IBD0 with each other, each parent will have transmitted both haplotype copies or all their genetic material. We applied this logic and compared the performance to that of using the expectation. The results of both approaches are similar, but the expectation yields slightly higher accuracy (not shown) and is also more computationally efficient to calculate. This is possibly due to uncertainty in the transmission status of each parent when the full siblings are all IBD1 with each other: at such locations we modeled the transmission rate of a given parent as the expectation of 75%. Another possibility is that false negative or false positive IBD segments adversely affect this estimation.

DRUID analyzes relatedness using full siblings and any of their half-siblings that it infers as equally related to *d*. It considers a set of half-siblings *H* (that are full siblings of one another; Figure S2B) to be equally related to *d* along with the primary full siblings in *S* if

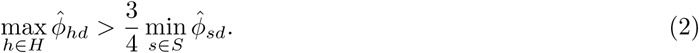

For the purposes of this analysis, given two or more sets of half-siblings, we take *S* to be the set containing the individual *s*\* with the highest overall 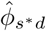 value. If the above equation does not hold, DRUID analyses *H* independently of *S*.

### Inferring IBD sharing of a grandparent using siblings and aunts/uncles

When data are available for a set of siblings together with some number of their aunts and uncles, the IBD segments these individuals share with a distant relative descend from a grandparent (Figure S2C). The expected proportion of a grandparent 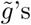 genome transmitted to these individuals is 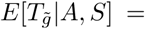 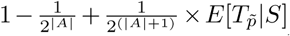 where *A* is the sets of aunts/uncles of the siblings in *S*, and 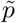 is the ungenotyped parent of *S*. Here, 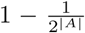 is the expected amount of the grandparent’s genome transmitted to his/her children in *A*, and the final term gives the expected amount of unique DNA transmitted to (|*A*| + 1)^st^ child 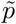 multiplied by the expected genome proportion that child transmitted to the grandchildren in *S*.

We estimate the proportion of the genome that the ungenotyped grandparent 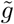 shares IBD with a distant relative *d* in an analogous way to that of ungenotyped parents:

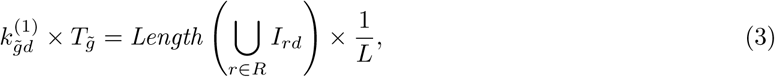

where *R* = *A* ∪ *S*. As above, we take the expectation of the left hand side and solve for 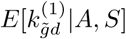.

As the siblings may have aunts/uncles both through their mother and their father, we group the aunts/uncles that are inferred to be siblings of one another to form up to two sets of aunts/uncles associated with a sibling set *S*, denoting these as *A*^(1)^ and *A*^(2)^. To infer relatedness to a distant relative *d*, DRUID includes a given set of aunts/uncles *A*^(*i*)^ of a sibling set *S* when

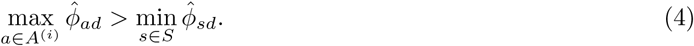

When this inequality holds for both sets *A*^(1)^ and *A*^(2)^, we use the set with the higher average 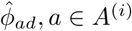. When no set fits this criteria, we continue the analysis using only the sibling set.

### Inferring IBD sharing between two sets of close relatives

In sufficiently large datasets or those with family-based recruitment, DRUID will infer several sets of closely related samples. When this occurs, distant relatedness may exist between two close relative sets and not merely to a single distant relative *d*. Inferring the amount of IBD shared between two ungenotyped ancestors from the two pedigrees enables inference at greater resolution than the potential alternative of using a single member of one of the pedigrees. Given two pedigrees 1 and 2 with corresponding sets *A*_1_, *A*_2_ of aunts/uncles and *S*_1_, *S*_2_ of siblings, we estimate the IBD sharing between two ungenotyped grandparents 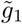 and 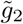 as

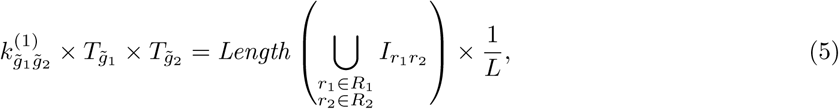

where *R_i_* = *A_i_*∪*S_i_* for *i* ∈ {1, 2}. Analogous equations apply for estimating relatedness between ungenotyped parents 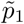 and 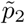 when both pedigrees only have sibling sets, and for cases where only one pedigree has an aunt/uncle set.

When IBD segments exist between at least one member of two sibling sets *S*_1_ and *S*_2_, DRUID performs this analysis. To determine whether to include available aunt/uncle sets, we consider the two pedigrees separately. For example, for pedigree 1, suppose siblings *S*_1_ have two aunt/uncle sets 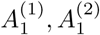 (through their two parents). Then, similar to Equation 4, we include a given set 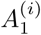 in the inference if:

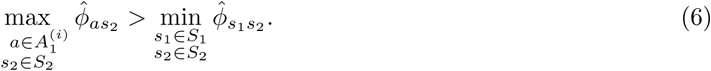

As before, if this inequality holds for both aunt/uncle sets, we include the set with higher average 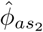 for 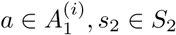. The inequality for pedigree 2 is analogous.

Given our assumptions, the only relatives that will share regions IBD2 with each other are full siblings. Therefore, distant relatives will not share IBD2 regions, but there are circumstances where their ungenotyped ancestors will have IBD2 sharing. In particular, given two sibling sets that are first cousins of each other, their parents are full siblings that consequently share non-trivial amounts of IBD2. It is important to account for any IBD2 when estimating the kinship coefficient between these ungenotyped ancestors.

Extending the IBD^(011)^ concept from inferring aunts/uncles to apply to first cousins, we infer regions that are IBD2 between the ungenotyped parents of two full sibling sets in the following way (Figure S3). First, for every pair of siblings *s*_1_, *s*_2_ ∈ *S*_1_, we locate regions in which they are IBD0 with each other. We then identify places these siblings are each IBD with two siblings from the other set *o*_1_, *o*_2_ ∈ *S*_2_. As long as this IBD inference is correct, these regions must be IBD2 in the parents of *S*_1_ and *S*_2_. Let 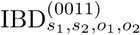 contain these regions for the indicated samples. Then, similar to Equation 5, we have

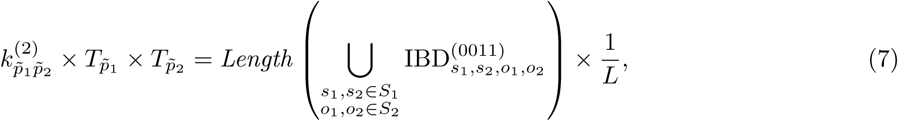

and we then use 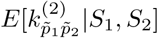 to compute 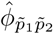 (adjusting for the fact that these IBD2 regions were also counted in the corresponding *k*^(1)^ term from Equation 5). DRUID performs this inference only when the initial *k*^(1)^-based estimate of 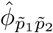 corresponds to a third degree or closer relationship. The above explanation considers two sets of siblings, but in general we apply this analysis using the oldest relevant generation of siblings available, which may be aunts/uncles.

### Determining degrees of relatedness for all sample pairs

To perform relatedness inference between all samples, DRUID must determine when to use standard pairwise relatedness measures (i.e., 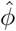 and 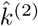 as first defined between two genotyped samples) and when to apply its multi-way inference. Prior to inferring the pedigree structures, DRUID infers a pairwise-only degree of relatedness between all pairs (Table 1). For pairs where neither is in a pedigree (connected component in the graph), DRUID reports this degree, which is the same as the pairwise Refined IBD estimate we compare to later. For pairs that are in the same pedigree structure, DRUID infers their degree of relatedness using this structure, reporting specific relationship types for first and second degree relatives when known (e.g., full siblings, avuncular). Otherwise, when one or both samples are in a (distinct) pedigree, the method determines whether each pedigree contains a parent or grandparent of a sample, and if so, whether that ancestor has the same or higher IBD sharing level with the other sample. If so, it successively moves up to older generations until arriving at the two oldest samples whose relatedness DRUID is to estimate.

Let *i, j* be the two samples with relatedness to be inferred where at least one is a member of a pedigree. If neither *i* nor *j* have any full siblings, half-siblings, aunts/uncles, or niece/nephews, DRUID reports the pairwise degree of relatedness between these samples. Alternatively, when one or both of *i, j* have full siblings, it uses these for inference, and also checks the relevant inequalities to determine whether to use any half-siblings, aunts/uncles, or niece/nephews. Following this, DRUID deduces the relatedness degrees between all the descendants of *i* and *j* in both pedigrees (setting each child as one degree further than its parent for all distant relatives).

When a pair of samples has a parent-child relationship but the parent is unknown, DRUID uses a heuristic to detect the parent. Specifically, suppose *i* and *j* have a parent-child relationship and DRUID needs to infer their relatedness to sample *d*. Without loss of generality, we consider *i* to be the parent of *j* if 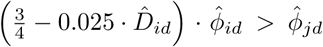, where 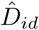 is the estimated degree of relatedness between *i* and *d*. This approach requires a larger relative difference between 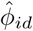 and 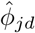 as the degree of relatedness between *i, j* and *d* increases. We have found this and the 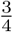 term (rather than the more intuitive 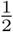) to be beneficial in both real and simulated data due to errors in the IBD detection. When this inequality does not hold, DRUID infers the degree of relatedness to *d* independently for *i* and *j*.

### Simulated relatives

We developed Ped-sim, a simulator for generating sample data from a specified pedigree structure that can have any number of generations and that is only (currently) constrained such that parents must marry either founders or samples from the same generation. Thus, Ped-sim is able to generate common relationship types including full and half-siblings, as well as less common types such as double cousins and various forms of inbred individuals. Ped-sim randomly samples founders from phased input samples and generates non-founders by simulating crossovers between the two haplotypes of all parents in the pedigree. The program uses either a sex-averaged genetic map or male- and female-specific genetic maps, with the sexes of parents randomized when using sex-specific maps. Additionally, Ped-sim can simulate crossovers either under a Poisson model or one that incorporates the effects of crossover interference^19,29,30^.

In the present study, we simulated pedigree samples using data from European-descent individuals collected from throughout Europe, the United States, Australia, and New Zealand^31^. We began by using all samples and SNPs listed (in files distributed by the European Genome-phenome Archive) as included by the original study following quality control procedures^31^. We further removed SNPs that failed a Hardy-Weinberg equilibrium filter (*P* < 10^−4^), had more than 5% missingness, or were outside the regions covered by the HapMap genetic map^32^, yielding a total of 10,299 individuals and 463,366 SNPs. Following phasing of all samples jointly with Beagle^33^ 4.1 (8 Jun 2017 release), we pruned SNPs for linkage disequilibrium (LD) using PLINK^34^ v1.90b2k with the command `–indep-pairwise 1000 25 0.25`, and then used PLINK to estimate pairwise relatedness with `–genome`. Next, we input these estimates to FastIndep^35^ to find a set of unrelated individuals. For purposes of filtering samples to use in this simulation, we considered all sample pairs with kinship 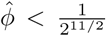, (i.e., fifth degree or more distant) as “unrelated.” Taking the resulting 8,955 phased samples as input, and using recently generated sex-specific genetic maps^36^ and a Poisson crossover model, we simulated pedigrees with Ped-sim v0.87.1b. Later, we simulated pedigrees that have only two full siblings, several parent-o spring pairs, and various second degree relationships (described below) using Ped-sim v0.93, this time incorporating crossover interference^19,29,30^. In both cases, Ped-sim introduced genotyping errors at a rate of 10^−3^ per site (with both alleles flipped at true homozygotes with 10% probability), and set a genotype to missing at a rate of 10^−3^ for the initial pedigrees and 5 × 10^−3^ for the secondary pedigrees. Because the sex-specific genetic maps cover a slightly smaller region than that spanned by the input SNPs, the simulated individuals included 462,828 SNPs.

The initial pedigree structures we simulated consist of distant relatives that are all some form of cousins. That is, their common ancestors are two grandparents from some number of generations ago depending on the degree of relatedness, as depicted in Figure S4. There are four types of these pedigrees: (1) five full siblings and one *D*^th^ degree relative, *D* ∈ {3, …, 10}; (2) two sets of five full siblings such that the sets are *D*^th^ degree relatives of one another, *D* ∈ {3, …, 10}; (3) five full siblings, two of their aunts/uncles, and one *D*^th^ degree relative, *D* ∈ {4, …, 10}; and (4) two sets that each contain five full siblings and two of their aunts/uncles such that the siblings in the two sets are *D*^th^ degree relatives of one another, *D* ∈ {5, …, 10}. As DRUID uses pairwise inference for first and second degree relatives, we chose *D* such that none of the distant relatives had relationships closer than third degree.

For these four pedigree types and each of their corresponding degrees of relatedness *D*, we simulated 120 replicate pedigrees, each randomized with respect to the assigned founder individuals and the sex of the parents. Specifically, we simulated the data in 10 batches, each batch containing 12 replicates of the various pedigree types and degrees. For any given run, Ped-sim assigns an input individual as a founder only once, but independent runs will in general assign some of the same input samples as founders. Thus, while the batches contain some overlapping founders, any specific pedigree structure is extremely likely to have different permutations of founders across batches.

To analyze the performance of aunt/uncle inference, we also simulated various forms of second degree relatives and included these in the above 10 simulation batches along with pedigree types 1–4. Specifically, each batch included: 12 sets of two full siblings and one aunt/uncle; 12 sets of two full siblings and one half-sibling; 6 sets of two full siblings and their two grandparents (with analysis done with respect each grandparent separately); and 12 sets of two full siblings and one double cousin.

To aid in the phasing step that precedes IBD detection, we incorporated an additional 1,674 unrelated samples in each batch, yielding a total of 5,022 individuals per batch. We used a Ped-sim option to output samples that were not used as founders in each batch, so the additional samples are drawn from the same European-descent sample data as the founders. Overall, one-third of each batch consists of these unrelated individuals (more precisely, fifth degree or more distant samples).

We further simulated a more complex pedigree structure shown in Figure S5, and we analyzed subsets of these samples in order to quantify the methods’ performance when utilizing various types of second degree relatives and/or a parent or child of target samples. These analyses focus on individuals L1, L2, or L3 (outline in red) and their *D*^th^ degree relationship to R1, for *D* ∈ {4, …, 10}. Here, we simulated 120 replicates for each degree *D* in three batches (40 replicates per batch). To aid phasing, we included an additional 1,540 unrelated samples per batch for a total of 5,460 samples in each batch.

We inferred IBD segments separately in each of the 13 simulation batches by running Refined IBD (part of Beagle 4.1) three times and reporting IBD regions as the union of identified segments across these runs (as recommended by the authors^26^). We used these segments in analyses of both the pairwise relatedness method we refer to as Refined IBD, and for running DRUID. Our analyses of pedigrees 1–4 range over different sizes of full sibling sets *S*, with |*S*| ∈ {2, 3, 4, 5}, where we randomly removed one sibling to obtain the set of siblings used for analysis of |*S*| − 1 siblings. Thus the included samples with smaller numbers of siblings are proper subsets of those with more. In a similar way, when analyzing the pedigrees that contain sets of aunts/uncles *A*, we tested with |*A*| ∈ {0, 1, 2}, and ensured that we include the same aunt/uncle for the range of sibling numbers |*S*| whenever |*A*| = 1. (When |*A*| ∈ {0, 2} the included aunts/uncles are necessarily identical for all values of |*S*|.)

PADRE^21^ requires input from PRIMUS^37^, a method for reconstructing pedigree structures using estimates of genome-wide IBD sharing of all pairs of samples. To infer these pedigrees, we ran PRIMUS v1.9.0 on each simulation batch (including the extra 1,674 unrelated samples), building pedigrees with all relationships up to second degree using the `–degree_rel_cutoff 2` option in two stages (the default of inferring up to third degree performed worse [not shown]). First, we used the `–no_IMUS` and `–no_PR` options so that PRIMUS only ran PLINK v1.90b2k to calculate genome-wide IBD estimates and did not perform pedigree reconstruction. We then subdivided the resulting `.genome` file (which lists PLINK’s genome-wide relatedness estimates for all pairs) into files containing only pairs of samples from each distinct pedigree structure (including the siblings, aunts/uncles, and all deep relatives). Afterwards we ran PRIMUS on each of these files, this time such that it inferred pedigrees, but with the subdivided files preventing PRIMUS from searching for relatives across distinct pedigree structures.

PADRE also requires initial likelihoods for pairwise degrees of relatedness from ERSA^38^, which itself requires IBD segments inferred by GERMLINE^39^. As part of the Refined IBD analysis noted above, we simultaneously inferred phase (i.e., using Beagle 4.1) in each batch, and then used GERMLINE version 1.5.1 (with options `-err_het 2 -err_hom 1 -min_m 1` as recommended by the ERSA authors^40^) to detect the IBD segments we provided to ERSA 2.0. Although PADRE’s documentation suggests using non-default parameters for ERSA, we find slightly poorer performance using those parameters as opposed to the defaults (not shown), and so we used the defaults. We then provided the ERSA results and the output from PRIMUS as input to PADRE. By default, PADRE infers only up to ninth degree relatives, and we modified this to enable it to infer up to degree 13. PADRE originally crashed when analyzing several pedigrees (22.5% of pedigree types 1–4 and 12.9% of the more complex scenarios), and we corrected these errors by removing error-generating PRIMUS pedigrees from consideration by PADRE. We applied this fix only when there were at least two alternate PRIMUS pedigrees for PADRE to consider. Following the fix, PADRE succeeded in analyzing 98.3% of type 1–4 pedigrees and 99.6% of the more complex scenarios. We excluded the results of all methods whenever PADRE crashed, which may give a slight advantage to PADRE. When successful, PADRE sometimes does not report a degree of relatedness between the distant relatives, and we treated this as a classification of unrelated.

### Real data

Besides simulated data, we evaluated DRUID and the other methods on real SNP array genotypes from the San Antonio Mexican American Family Studies (SAMAFS)^41–43^. Specifically, we analyzed 2,485 individuals typed at 521,184 SNPs from one of several dozen pedigrees. We previously described our quality control procedures for these data^23^, but in brief, we set sites with Mendelian errors to missing and mapped the SNP probes to GRCh37. Following SNP filters based on dbSNP and other auxiliary databases, we filtered SNPs with more than 2% missingness and samples with more than 10% missingness. The samples were typed on several Illumina SNP arrays, and our filters resulted in the inclusion of SNPs that are common to all arrays. We also excluded from consideration 2,618 pairs of samples (out of more than 3 million) that have evidence of being descended from parents that are cryptically related.

To analyze these data, we phased and detected IBD in all 2,485 individuals by running Refined IBD three times (phase generated simultaneously using Beagle 4.1), taking the union of the IBD segments identified in all runs. As in the simulated data, we ran PRIMUS (incorporating first and second degree relationships into the pedigrees) in two stages, first to obtain genome-wide IBD sharing estimates, and second on subdivided (PLINK generated) `.genome` files. These subdivided files restrict PRIMUS to analyzing two sets of close relatives to be analyzed (described next). We also ran GERMLINE on the phased data for all 2,485 samples followed by ERSA, both using recommended settings. Finally, we ran PADRE using the output of PRIMUS and ERSA and with the modification to enable it to infer up to relatedness degree 13. Similar to the simulations, PADRE originally crashed in a subset (5.2%) of these SAMAFS analyses. We corrected these errors by removing error-generating PRIMUS pedigrees when doing so left at least two alternate pedigrees for PADRE to consider. With this fix, PADRE successfully analyzed 97.4% of SAMAFS pedigrees, and we again excluded results from both DRUID and PADRE in any cases of PADRE failure.

We analyzed the performance of DRUID and PADRE on the SAMAFS data, leveraging the pedigree-based recruitment of this sample which includes reported pedigree structures (example in Figure S6). Specifically, we analyzed two reported close relative sets that are also reported to be distantly related to one another. Any misreported relationships have the potential to confound this analysis, though they should equally affect both DRUID and PADRE and in general be fairly infrequent. We analyzed two distantly related sets of full siblings, and when one or both sibling sets had associated aunts/uncles available, we also analyzed the same samples after adding one (choosing randomly if there are two) and then both sets of aunts/uncles.

To evaluate the performance of DRUID’s aunt/uncle inference procedure, we considered all reported sets of ≥ 2 full siblings and their reported second degree relatives. Once again, misreported relationships may confound this analysis, although our results suggest most relationships are correct. We randomly selected two siblings from each set of full siblings for analysis, repeatedly sampling non-overlapping pairs of siblings when more than two siblings are available. Then, for every selected pair of siblings, we analyzed their *k*^(2)^ and IBD^(011)^ levels to all their reported second degree relatives. Note that whenever multiple pairs of siblings from the same sibling set exist, this considers the same second degree relatives several times.

As a more conservative analysis, we also ran DRUID and Refined IBD on a restricted set of samples consisting of confidently inferred sets of full siblings and their aunts/uncles together with only one distant relative. For this, we filtered reported full siblings to those whose inferred pairwise degree of relatedness is one, obtaining sets in which all pairs of siblings have a first degree relationship. To ensure the reported aunts/uncles of these full sibling sets are correct, we checked whether the pairwise relationship of a given aunt/uncle with each full sibling is second degree. If this is case, we accepted this individual as an aunt/uncle. Because inference of second degree relatives has a slightly lower power than first degree inference^23^, for each aunt/uncle verified in this manner, we also include any confident full siblings (based on the first degree relationship criteria) of this person as aunts/uncles. We also performed an analysis using half-siblings, and for this purpose, we generated confident sets of half-siblings in the same manner as for aunts/uncles. To increase sample size, we analyzed the inference rates using each set of close relatives and all their available distant relatives of a specified degree (the latter analyzed one at a time). Therefore, results include multiple analyses of the same close relative sets for various distant relative, treating these as independent.

## Results

To evaluate DRUID, we examined the rate at which it correctly classifies the degree of relatedness—both exactly and to within one degree of the truth—between simulated and real samples compared to PADRE and Refined IBD. The latter approach uses the IBD regions detected by Refined IBD to infer a pairwise degree of relatedness. This is the same approach DRUID employs to infer first and second degree relatives and individuals not in pedigrees and is among the top performing pairwise relatedness methods^23^.

PADRE uses a composite likelihood method to infer relatedness between two networks of samples, with each network containing individuals whose relationship to one another must be represented in an input pedigree that PRIMUS inferred. In the presence of ambiguous relationships, PRIMUS outputs multiple possible structures together with their likelihoods and PADRE considers each of these likelihoods and their corresponding structures in its analysis. PADRE also takes as input pairwise relatedness likelihoods inferred by ERSA and uses these to identify the degrees of relatedness that maximize the overall composite likelihood. The full composite likelihood for a given pair of pedigree structures and relatedness degrees between distant relatives is the product of the pedigree likelihoods (from PRIMUS) and all pairwise relationship likelihoods (from ERSA) that exist across the two networks. Because PADRE is intended to apply to two non-singleton networks of samples, we ran it on the simulated pedigree structures that contain two non-singleton close relative sets (i.e., types 2, 4, and those with various second degree relationships). Results from running DRUID and Refined IBD on pedigree types 1 and 3 are in Figures S7, S8, S11, and S12. For relatedness inference of singletons (i.e., traditional pairwise inference) our recent analysis showed that ERSA and Refined IBD (used by DRUID) have comparable performance^23^. This similarity of ERSA and Refined IBD indicates that the below comparisons of PADRE and DRUID are informative about these latter methodologies and not differences in the IBD detection methods they each use.

Below we briefly describe DRUID’s performance for inferring aunts/uncles and then proceed to compare it to Refined IBD and PADRE for distant relatedness inference in simulated and real data. For the distant relatedness inference that includes varying numbers of siblings and optional aunts/uncles, we report classification rates only for pairs in the sibling sets (i.e., excluding the aunts/uncles). This enables a direct comparison of results for the same degree of relatedness with and without aunt/uncle inclusion. For simulated pedigree types 1 and 3, which have only one distant relative, the target number of pairs of distantly related samples to classify is |*S*|. For pedigree types 2 and 4, there are |*S*|^2^ pairs to classify. For the simulations that are enriched in second degree relatives, we analyze classification rates for only one pair of samples: R1 and either L1, L2, or L3 (Figure S5). We compute the classification rate for a given collection of distant relatives by averaging over the indicated number of pairs; we then average these rates over all distant relative collections (i.e., over each instance of two distantly related sets of close relatives). In simulations, this average is over the 120 replicate structures, and in real data, we report the numbers of analyzed structures.

### Inferring aunts and uncles with DRUID

We used the real SAMAFS data as well as simulated individuals to examine DRUID’s performance for aunt/uncle classification. As noted earlier, in nearly all cases, a pair of siblings *s*_1_ and *s*_1_ and one aunt/uncle *a* have more than 50 cM of the 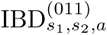 pattern DRUID uses for classification (Figures 2B and S1B). Indeed, considering SAMAFS data for two reported full siblings and one reported second degree relative, DRUID infers 92.2% of all aunt/uncles as such with zero false positives (total 1449 aunts/uncles out of 1893 second degree relatives). Results using simulated data are similar: with two full siblings and various second degree relatives (Methods), DRUID recovers 90.1% of aunts/uncles, also with zero false positives (total 120 aunts/uncles out of 480 second degree relatives). The filter that removes double cousins from consideration (Figure 2A and S1A) is highly effective as it captures all double cousins in both real and simulated data with zero false positives (total 38 real and 120 simulated double cousins).

### Distant relative inference using siblings and aunts/uncles

Figure 3 depicts results for type 2 pedigrees (i.e., those with two distantly related sets of siblings), subdivided into exact and within-one-degree inference accuracy for |*S*| = 2 and |*S*| = 5, and for most degrees of relatedness *D* (full results in Figures S9 and S10). When analyzing type 2 pedigrees, both DRUID and PADRE outperform the pairwise method Refined IBD for *D* = 3 to 9, with between 5.0–35.4% more relatives classified exactly for these degrees (Figure 3A). DRUID and PADRE show greater improvement relative to Refined IBD when |*S*| = 5 than |*S*| = 2, consistent with the methods having more information with which to make inference for larger |*S*|. The differences among these methods are more modest for *D* = 10, presumably because there is limited information to perform classification at this degree. Indeed, DRUID only classifies 28.3% of relatives exactly for *D* = 10. Despite this, DRUID’s within-one-degree classification accuracy is quite high for *D* = 10 at 79.6–80.0% (Figure 3B).

**Figure 3:**
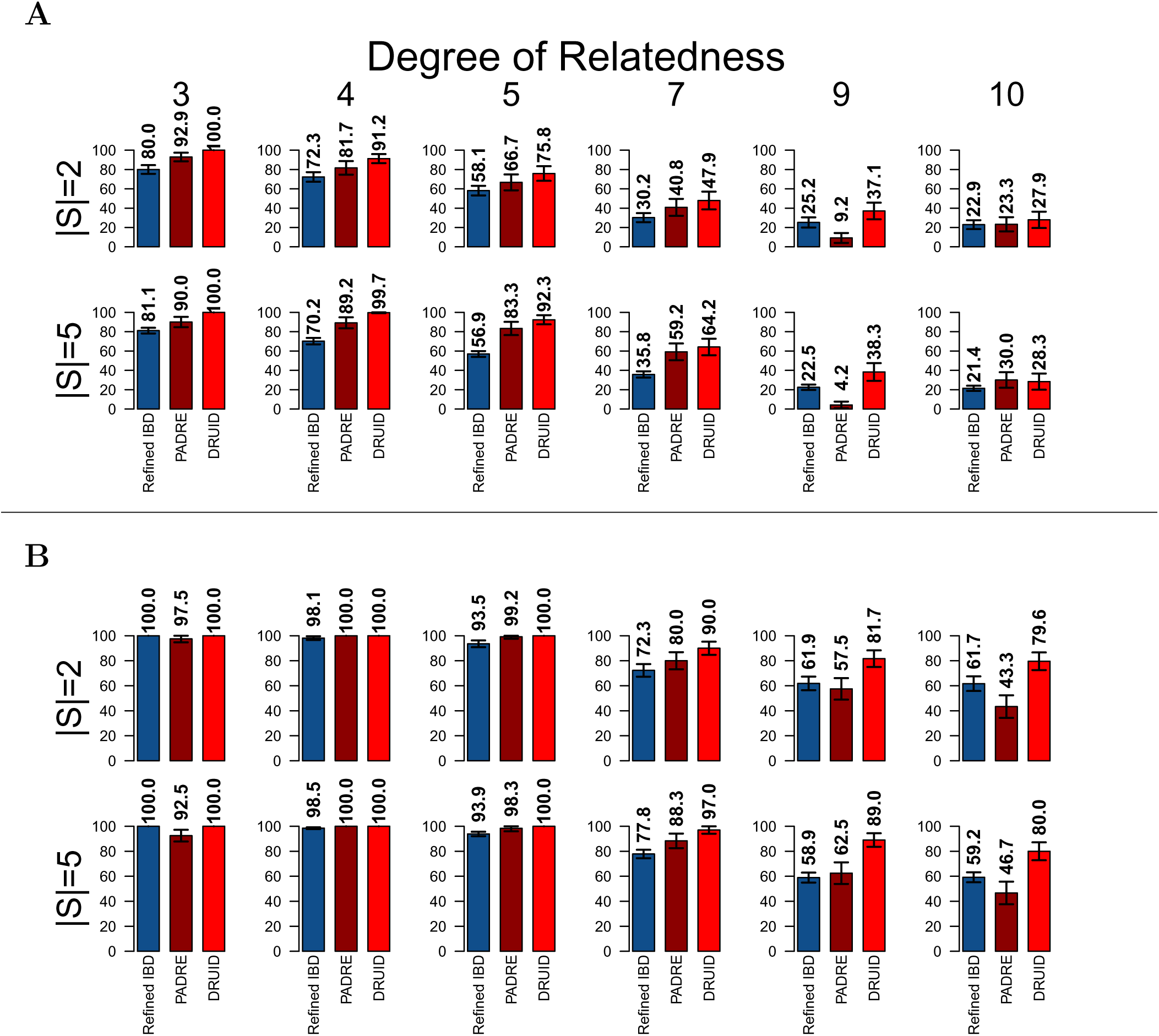
(**A**) For simulated type 2 pedigrees, average percent of distantly related sample pairs from the two sibling sets that are inferred as their true degree of relatedness using Refined IBD, PADRE, and DRUID. Raw averages are listed above each bar. Rows of bar plots have the same number of siblings included in both sibling sets, indicated as |*S*| (left). Columns show results for different degrees of relatedness, with the true degree listed above. (**B**) As in (A), but shows the average percent of distant relatives inferred to be related as the true degree *D* or as *D* ± 1. Error bars are bootstrapped (over complete relative sets) 95% confidence intervals.

For the multi-way relatedness methods, DRUID shows greater accuracy than PADRE for most values of *D*, classifying up to 10.5% more pairs exactly for *D* = 3 to 7 (Figure 3A). Considering within-one-degree classification, DRUID has 100% accuracy up to *D* = 5, with PADRE showing some (though modest) inaccuracies up to this degree (Figure 3B). However, for more distant degrees, PADRE’s within-one-degree classification rates drop relative to DRUID. For example, when |*S*| = 5, PADRE infers only 62.5% and 46.7% of sample pairs within-one-degree for *D* = 9 and *D* = 10, respectively. By contrast, DRUID infers, respectively, 89% and 80% of these relatives to within one degree of the truth. For exact inference of higher relatedness degrees, PADRE’s accuracy drops when *D* = 9, but is more competitive with DRUID when *D* = 10. As noted earlier, by default PADRE only infers up to ninth degree relatives, and its within-one-degree accuracies are similar when using this default compared to these results (not shown). However, with those settings, it infers 0% of tenth degree relatives exactly. Thus, while there is an apparent bias against ninth degree inference in the settings we used, the alternative always misclassifies tenth degree relatives.

The differences between the methods are even more striking when inferring relatedness within type 4 pedigrees, which includes siblings and up to two aunts/uncles, as shown in Figure 4 (full results in Figure S13 and S14). For *D* = 5 and *D* = 7, when analyzing data for all |*A*| = 2 aunts/uncles and |*S*| = 5 siblings (from both close relative sets), DRUID infers 23.3% and 31.3% more pairs exactly correct, respectively (Figure 4A). Considering higher relatedness degrees, the downward bias in PADRE for *D* = 9 remains, resulting in a sizable discrepancy between the two methods. When *D* = 10, and again leveraging all data (|*A*| = 2, |*S*| = 5), DRUID infers 10.7% more relatives exactly than PADRE.

**Figure 4:**
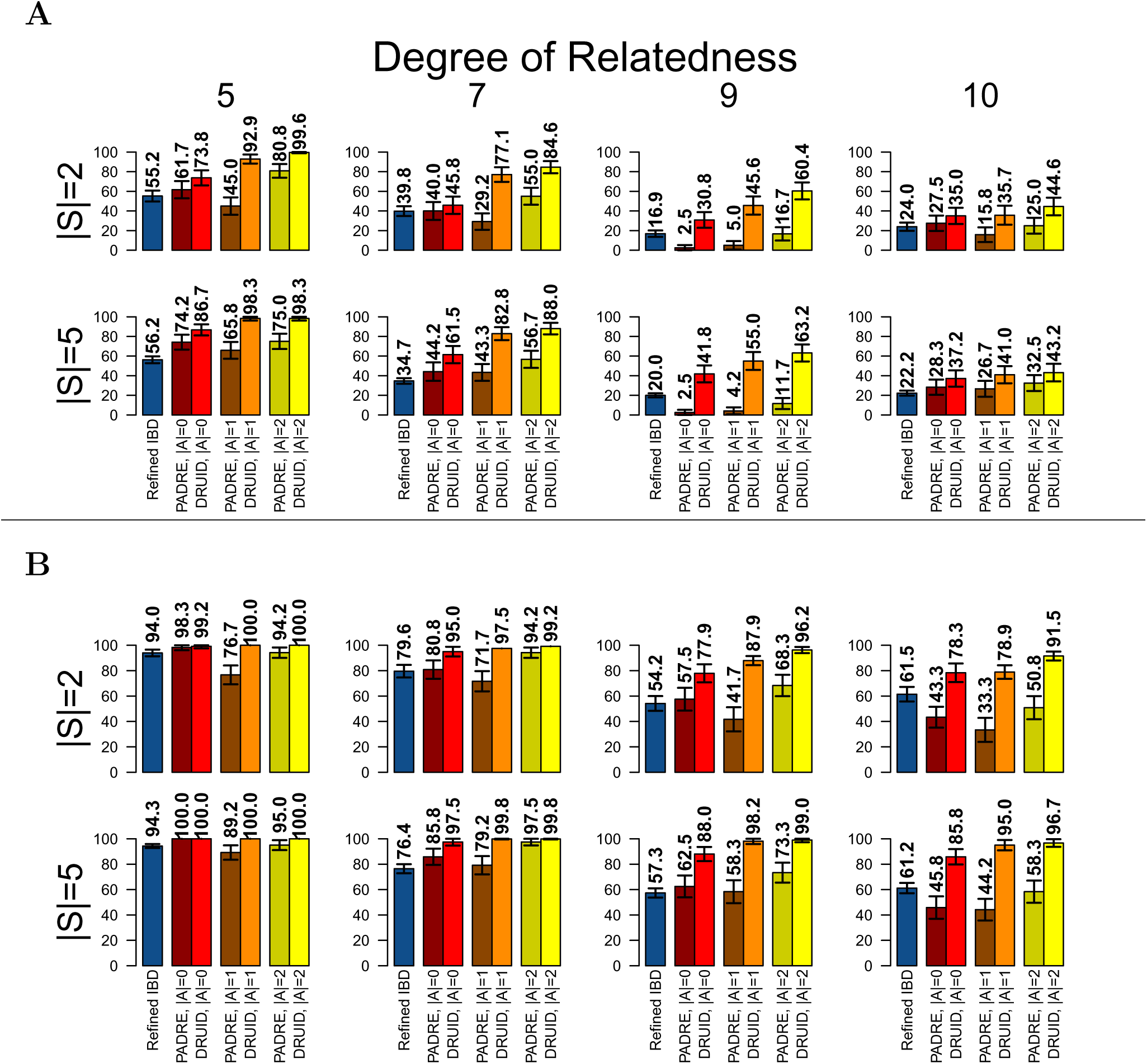
(**A**) For simulated type 4 pedigrees, average percent of distantly related sample pairs from the two sibling sets (bottom generation in Figure S4) that are inferred as their true degree of relatedness using Refined IBD, PADRE, and DRUID. Raw averages are listed above each bar. Rows of bar plots have the same number of siblings included in both sibling sets, indicated as |*S*| (left). Analyses with different numbers of aunts/uncles |*A*| included are shown in distinct bars as labeled. Columns show results for different degrees of relatedness between the two sibling sets, with the true degree listed above. Analyses with |*A*| = 0 parallel those for type 2 pedigrees and use siblings only from the type 4 pedigree simulations. (**B**) As in (A), but shows the average percent of distantly related siblings inferred to be related as the true degree *D* or as *D* ± 1. Error bars are bootstrapped (over complete relative sets) 95% confidence intervals.

One of the key reasons DRUID is likely to outperform PADRE is its ability to detect the aunts/uncles correctly and not consider them as half-siblings or grandparents. The effect of the ambiguity in second degree relatives is especially evident when |*A*| = 1: in most instances, PADRE classifies fewer relatives correctly for |*A*| = 1 than when |*A*| = 0 (i.e., with less data). When |*A*| = 1, PADRE commonly infers the aunt/uncle as a grandparent or half-sibling (Figure S15), which sometimes leads it to classify the siblings one degree differently than it otherwise would. This effect is diminished for |*A*| = 2 (Figure S15) because PRIMUS readily detects these aunts/uncles as full siblings of one another and benefits from the fact that a grandparents’ sibling should be one degree more distantly related to the siblings, whereas aunts/uncles (and half-siblings) should be equally related. Overall, DRUID’s aunt/uncle inference is effective and appears to enable sizable gains in inferring relatives to their exact relatedness degree and within one degree of this.

The within-one-degree inference results for type 4 pedigrees show similar patterns to those from the exact inference (Figure 4B). Considering *D* = 5 and *D* = 7, and when |*A*| ≥ 1, DRUID infers between 99.2–100% of all pairs correctly or different by one degree (Figures 4B, S14). PADRE is competitive with DRUID for these two degrees when |*A*| = 2, but underperforms for |*A*| = 1. When *D* = 9 and *D* = 10, and while |*A*| = 2 and |*S*| = 5, DRUID classifies 96.7–99.0% of pairs to within one degree of the truth; at the same time, PADRE infers 58.3–73.3% of these relatives correct or off by one degree. Thus DRUID is highly accurate at inferring deep relationships to within one relatedness degree for pedigrees containing siblings and their aunts/uncles.

### Distant relative inference using a wider range of close relationships

Because of challenges in distinguishing second degree relatives other than aunts/uncles of two or more siblings, DRUID does not leverage half-siblings, grandparents, or undetected avuncular relatives (unless specified in the input). However, PRIMUS and therefore PADRE consider all possible pedigrees consistent with detected second degree relatives and may consequently be better than DRUID in settings where data for full siblings are lacking. To evaluate such scenarios, we analyzed subsets of individuals from the simulated pedigree structure in Figure S5. In each of the scenarios, because PADRE requires distant relatives to be non-singletons, one side of the pedigree has two cases: a parent-child pair or a pair of full siblings. More specifically, the analyses always include (labels from Figure S5) sample R1, either R2 or R3 (R1’s child or sibling, respectively), and a sample *i* ∈ {L1, L2, L3} together with: one second degree relative of *i* (three scenarios); one second degree relative *r* of *i* and the child of *r* (three scenarios); two second degree relatives of *i* (two scenarios); the child of *i* and either no other samples or one second degree relative of *i* (four scenarios); and the parent of *i* and either no other samples or one second degree relative (three scenarios). We report classification rates for each of these 15 scenarios as that for the relationship between R1 and *i* only, defining *D* below as the degree of relatedness between these samples.

Figures S16 and S17 depict the classification rates for Refined IBD, PADRE, and DRUID in all 15 scenarios and the two cases (R1 and child or R1 and sibling), averaged over all 120 replicates. Considering *D* ∈ {4, 5, 6}, the results are mixed, with DRUID outperforming PADRE in 56 of 84 of these settings, PADRE beating DRUID in 24, and with equal performance in four settings. (Note that when *D* = 4, the grandparent [GP] of *i* ∈ {L1, L2, L3} is a second degree relative of R1 and thus PRIMUS detects all samples as part of the same pedigree. PADRE is therefore not applicable in the 6 settings where *D* = 4 and the GP is included.) In all but one scenario, PADRE classifies up to 6.7% more pairs correctly than DRUID, and DRUID infers up to 18% more. The outlying scenario arises when using L1, L1’s aunt/uncle (AU) and L1’s GP: here, PADRE infers 3.3–30% more pairs correctly than DRUID up to *D* = 7. In this scenario, PRIMUS is effective at limiting the scope of possible relationships among the samples, yielding considerably higher performance than DRUID. Turning to deeper relationships, when *D* ≥ 7 (*D* ≥ 8 for the aforenoted L1 with GP and AU scenario), DRUID outperforms PADRE in all settings, classifying 5.8–38% more distant relatives to their correct degree. Note that whenever parent-child and full sibling relationships are lacking, DRUID classifies relatives in the same way as Refined IBD. Given data for the parent of a target sample, and with no full sibling data, DRUID’s heuristic parent detection often improves performance, with up to 26.7% more samples correctly detected compared to Refined IBD.

We also evaluated PADRE’s second degree relationship type inferences for L1’s GP, AU, and HS relatives (Figure S18). As might be expected, PADRE is most able to determine these relationships for smaller *D*. In general, PADRE appears to be somewhat biased against inferring GP relationships, with some favor for HS classification. Note that PADRE’s classification rates depend on the number of first and second degree relatives available—as well as the pedigree structure of the distant relatives—and these results are specific to scenarios that include only L1 and one second degree relative and R1 and a child or full sibling.

Next, we examined the accuracy of PADRE and DRUID when classifying relatives in these scenarios to within one degree of the truth. Under this metric, excluding the L1 with GP and AU scenario, PADRE and DRUID have much closer performance for *D* ∈ {4, 5}, with the methods always within 3.3% of each other (not shown). For *D* = 6, PADRE performs well, classifying up to 13.3% more samples to within one degree in some scenarios while DRUID classifies up to 3.3% more (not shown). However, for *D* ≥ 7 (*D* ≥ 9 for L1 with GP and AU), DRUID always has equal or improved performance relative to PADRE, inferring up to 71.7% more relatives correct or off-by-one degree (average of 31.4% more for these scenarios; not shown).

In summary, PADRE is extremely accurate in the L1 with GP and AU scenario, while DRUID is otherwise either comparable to PADRE (except when *D* = 6 for off-by-one classification) or, for *D* ≥ 7 has notably higher classification rates.

### Computational complexity and runtime

To analyze DRUID’s runtime in a range of sample sizes, we used random subsets of the 5,460 samples from each batch of the simulations that are enriched in second degree relatives. We found that DRUID scales quadratically in these analyses with an average runtime (over the three batches) of 24.6 minutes for 288 samples; 4.4 hours for 577 samples; and 44.8 hours for 1,155 samples (Table S1). In all these runs, DRUID used no more than 5.5 GB of RAM. We note that DRUID is parallelizable, and given these times, we believe it is capable of analyzing several thousand samples simultaneously. PADRE is faster than DRUID as it completes the analysis of 1,155 samples in an average of 1.9 hours. However, PADRE requires substantially more memory, using an average of 43.6 GB to analyze 1,155 samples (Table S1).

A bottleneck in any analysis of IBD segments is the initial detection of those segments. For example, to analyze the batches with 5,460 samples, Refined IBD took an average of 17.3 days (wall clock time) when run with 12 threads. Additionally, (as recommended) we ran it three times for each batch. Therefore, scaling to datasets with more than roughly 20,000 samples may require alternate strategies, such as using Eagle^44^ to phase followed by GERMLINE to detect IBD.

### Distant relative inference in SAMAFS data

Our initial distant relative analysis in the SAMAFS data focused on the performance of DRUID in comparison to Refined IBD. When using sets of confidently inferred close relatives (Methods) consisting of either one set of full siblings and a distant relative (Figure S19), or siblings, some number of aunts/uncles, and a distant relative (Figure S20), the classification results largely recapitulate those from simulations. In particular, including larger numbers of close relatives leads to greater accuracy gains by DRUID, with inference using aunts/uncles providing dramatic improvement relative to Refined IBD. One distinct feature of the results in these real data compared to simulations is a general increase in accuracy rates for both methods. This may indicate that the simulation results underestimate the accuracy of all the methods, which would be expected if the simulated individuals have higher variance in IBD sharing than occurs in real data. Alternatively, because the SAMAFS data contain a relatively small number of pedigrees, each with many close relatives, the quality of phasing (and therefore of detected IBD segments) may be higher than in the simulated data.

Following this analysis, we used the SAMAFS data to analyze whether including confidently inferred half-siblings provides the same classification rates as including only full siblings. As shown in Figure S21, the accuracy rates are effectively identical when using two or three full siblings as when using one half-sibling together with one or two full siblings. This is as expected since IBD detection between distant relatives should not be heavily influenced by the exact types of close relationships that exist among other samples.

Finally, we compared PADRE to DRUID using relative sets from SAMAFS, relying only upon reported relationships whether these fit the confident inference criteria or not. We performed the inferences with two sets of full siblings that are distantly related to each other and did not restrict the size of the sibling sets; these range from |*S*| = 2 to 11. Where available, we also included one or two sets of aunts/uncles that are associated with one or both of the sibling sets, denoting the number of included aunt/uncle sets as 𝒜. We required all included aunt/uncle sets to have size |*A_i_*| = 2, randomly selecting two samples when more than two are available. To increase the sample sizes, we performed inference with the maximum 𝒜 and also randomly removed aunt/uncle sets so that we included all pairs of sibling sets that have the needed data for each 𝒜. (Thus included samples with smaller 𝒜 for a given degree are a superset of those with larger 𝒜.) As in earlier analyses, the reported classification rates are only for distant relatives in the two sibling sets.

Figure 5 depicts the results of this analysis. The rate of classifying relatives to their exact relatedness degree is higher in DRUID than in PADRE for degrees *D* ≥ 4, ranging between 1.3–9.2% more correct inferences using DRUID (Figure 5A). For *D* = 3, PADRE’s exact classification rates exceed those of DRUID by 0.6%, but a caveat is that PADRE crashed in 13.2% of the *D* = 3 analyses and we did not count these cases for either method. Notably DRUID inferred all relationships correctly as *D* = 3 in the omitted analyses. As observed in the simulated data, DRUID outperforms PADRE by a larger margin when using aunts/uncles than using only siblings, though both methods generally improve when given aunt/uncle data. Both methods achieve 100% classification rates when inferring relatives to within one degree of their true relationship (Figure 5B), which is generally consistent with the simulation results for these degrees. Overall, this analysis in SAMAFS demonstrates the effectiveness of performing inference based on the reconstructed IBD sharing profile of an ungenotyped ancestor, as DRUID’s accuracy results typically exceed both pairwise and another multi-way inference method.

**Figure 5:**
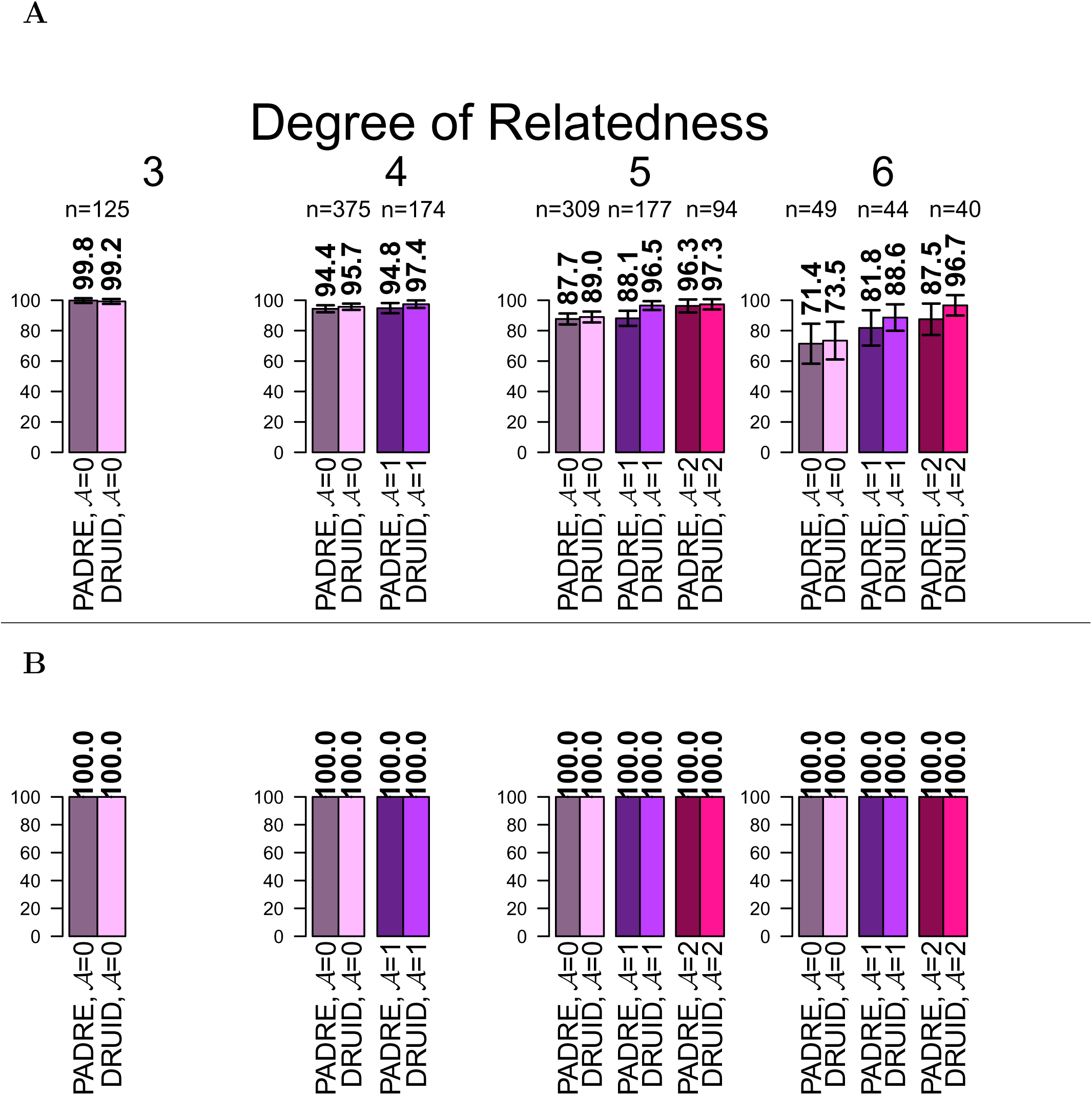
(**A**) Rates of DRUID and PADRE inferring a range of degrees of relatedness between two full sibling sets in SAMAFS that are distantly related to each other. Raw average inference rates are listed above each bar. Analyses consider inclusion of two full sibling sets only (𝒜 = 0), two sibling sets and one associated aunt/uncle set (𝒜 = 1), and two sibling sets that each have an aunt/uncle set (𝒜 = 2), as indicated by bar labels. Number *n* of analyzed collections of two close relatives sets shown above. (**B**) As in (A), but shows the average percent of distant relatives inferred to be related as the true degree *D* or as *D* ± 1. Error bars are bootstrapped (over complete relative sets) 95% confidence intervals.

## Discussion

Classifying relatedness between samples is a classical problem in genetics^9^, and is relevant to both trait association mapping^8,11,12^ as well as population genetics^1^. Recent advances in this domain have come through the analysis of IBD segments^40^ instead of only independent markers, and through analyses that leverage signals from multiple individuals^21,22^ instead of only pairs.

We developed and evaluated DRUID, an algorithm that uses IBD sharing between multiple close relatives in order to infer the IBD sharing profile of one of their ungenotyped ancestors. A critically important factor in DRUID’s performance is the correct inference of the pedigree relationships among the close relatives. This is so because mistakes in these relationships lead to reconstruction of IBD sharing either of an incorrect individual or one that does not exist—e.g., composed of a combination of IBD segments from a grandparent and a parent. Because the relatedness inferred for the genotyped individuals derive from these unobserved samples, mistakes have the potential to bias the results. To avoid such issues, we focus our inference on full siblings and we developed a new approach to accurately detect aunts and uncles of a pair of siblings. The aunt/uncle inference quality is very high with zero false positives and 92.2% and 90.1% recall in real and simulated data, respectively. DRUID can also use directly observed ancestors such as parents and specified grandparents to aid inference.

Our analyses show that DRUID and PADRE both perform substantially better than Refined IBD, with up to 35.4% more relatives classified correctly in simulations when data for two distantly related sets of full siblings are available (Figure 3A). Considering DRUID and PADRE only, for relatedness between two sets of full siblings, DRUID infers up to 10.5% more pairs correctly in simulations (Figure 3A), and up to 2.1% more relatives correctly in SAMAFS (Figure 5A). In turn, for data comprising two sets of full siblings with up to two aunts/uncles, DRUID infers up to 10.7–31.3% more simulated pairs to the correct degree of relatedness compared to PADRE (Figure 4A), and up to 9.2% more relatives in SAMAFS (Figure 5B). This dramatic improvement stems from DRUID’s approach in general as well as from its aunt/uncle inference.

Ideally, methods would be able to infer the degree of relatedness between samples to their exact values—and DRUID makes possible such specific inference for fifth degree degree relatives given data for two sets of full siblings (75.8–92.3% exactly correct in simulations, Figure 4A; 89.0% exactly correct in SAMAFS, Figure 5A). Still, inference to within one degree of the true relationship is very informative, and DRUID infers between 79.6–96.7% of simulated tenth degree relatives in this way (Figures 3B and 4B). This high rate of classification for such distant relatives signals a sizable shift in the effectiveness of such inference. Because the coefficient of variation in IBD sharing increases for more distant relatedness, exact inference is most likely to only be achievable for lower relatedness degrees. Yet exact inference isn’t required in all settings, and identifying relatedness to within a range of degrees provides information that is useful for further characterization. One noteworthy application area is that of genetic genealogy, a service offered to consumers by several genetic testing companies, and which has become especially popular in recent years as testing prices have fallen.

Going forward, a possible extension of this work is to effectively iterate DRUID’s approach: identify and classify the relationships of two sets of distant relatives, and then use that classification to reconstruct the IBD sharing profile between their mutual ancestor(s) and other even more distant relatives. Success in this context will require either modeling uncertainty in which ancestor is being reconstructed, or alternative methodologies that enable more precise knowledge of the pedigree structure that exists between the initial two sets of close relatives. Another tantalizing possibility in this domain is that of directly inferring genetic data (such as haplotypes) for specific ancestors. While DRUID infers information about ancestors, it is ambiguous about which ancestor is being analyzed. Nevertheless, its approach is effective and opens the door to more detailed pedigree-based analyses in studies that contain many relatives—now including nearly all newly generated datasets.

## Web resources

DRUID:http://github.com/williamslab/druid/

Ped-sim: http://github.com/williamslab/ped-sim/

## Acknowledgements

We thank the anonymous reviewers for helpful suggestions. This work was supported by a National Science Foundation Graduate Research Fellowship grant number DGE-1144153 to M.D.R.; Qatar National Research Fund grant NPRP 7-1425-3-370 to J.G.M.; an Alfred P. Sloan Research Fellowship, and a seed grant from Nancy and Peter Meinig to A.L.W. The SAMAFS are supported by NIH grants R01 HL0113323, P01 HL045222, R01 DK047482, and R01 DK053889. This study makes use of data generated by the Wellcome Trust Case-Control Consortium. A full list of the investigators who contributed to the generation of the data is available from www.wtccc.org.uk. Funding for the project was provided by the Wellcome Trust under award 076113, 085475 and 090355

## References

[1] John Wakeley, Léandra King, Bobbi S Low, and Sohini Ramachandran. Gene genealogies within a fixed pedigree, and the robustness of kingmans coalescent. Genetics, 190(4):1433–1445, 2012.

[2] Clare Bycroft, Colin Freeman, Desislava Petkova, Gavin Band, Lloyd T Elliott, Kevin Sharp, Allan Motyer, Damjan Vukcevic, Olivier Delaneau, Jared O’Connell, et al. Genome-wide genetic data on 500,000 UK Biobank participants. bioRxiv, page 166298, 2017.

[3] Monkol Lek, Konrad J Karczewski, Eric V Minikel, Kaitlin E Samocha, Eric Banks, Timothy Fennell, Anne H O’Donnell-Luria, James S Ware, Andrew J Hill, Beryl B Cummings, et al. Analysis of protein-coding genetic variation in 60,706 humans. Nature, 536(7616):285–291, 2016.

[4] Frederick E Dewey, Michael F Murray, John D Overton, Lukas Habegger, Joseph B Leader, Samantha N Fetterolf, Colm O’Dushlaine, Cristopher V Van Hout, Jeffrey Staples, Claudia Gonzaga-Jauregui, et al. Distribution and clinical impact of functional variants in 50,726 whole-exome sequences from the DiscovEHR study. Science, 354(6319):aaf6814, 2016.

[5] Jeffrey Staples, Evan K Maxwell, Nehal Gosalia, Claudia Gonzaga-Jauregui, Christopher Snyder, Alicia Hawes, John Penn, Ricardo Ulloa, Xiadong Bai, Alexander E Lopez, Cristopher V Van Hout, Colm O’Dushlaine, Tanya M Teslovich, Shane E McCarthy, Suganthi Balasubramanian, H Lester Kirchner, Joseph B Leader, Michael F Murray, David H Ledbetter, Alan R Shuldiner, George Yancoupolos, Frederick E Dewey, David J Carey, John D Overton, Aris Baras, Lukas Habegger, and Jeffrey G Reid. Profiling and leveraging relatedness in a precision medicine cohort of 92,455 exomes. bioRxiv, 2017.

[6] Oriol Canela-Xandri, Konrad Rawlik, and Albert Tenesa. An atlas of genetic associations in UK Biobank. bioRxiv, page 176834, 2017.

[7] Eunjung Han, Peter Carbonetto, Ross E Curtis, Yong Wang, Julie M Granka, Jake Byrnes, Keith Noto, Amir R Kermany, Natalie M Myres, Mathew J Barber, et al. Clustering of 770,000 genomes reveals post-colonial population structure of north america. Nature Communications, 8:14238, 2017.

[8] Benjamin F Voight and Jonathan K Pritchard. Confounding from cryptic relatedness in case-control association studies. PLOS Genetics, 1(3):e32, 2005.

[9] Bruce S Weir, Amy D Anderson, and Amanda B Hepler. Genetic relatedness analysis: modern data and new challenges. Nature Reviews Genetics, 7(10):771–780, 2006.

[10] Joshua G Schraiber and Joshua M Akey. Methods and models for unravelling human evolutionary history. Nature Reviews Genetics, 16(12):727–740, 2015.

[11] Hyun Min Kang, Jae Hoon Sul, Noah A Zaitlen, Sit-yee Kong, Nelson B Freimer, Chiara Sabatti, and Eleazar Eskin. Variance component model to account for sample structure in genome-wide association studies. Nature genetics, 42(4):348–354, 2010.

[12] Xiang Zhou and Matthew Stephens. Genome-wide efficient mixed-model analysis for association studies. Nature genetics, 44(7):821–824, 2012.

[13] Sharon R Browning and Brian L Browning. Haplotype phasing: existing methods and new developments. Nature Reviews Genetics, 12(10):703–714, 2011.

[14] Catarina D. Campbell, Jessica X. Chong, Maika Malig, Arthur Ko, Beth L. Dumont, Lide Han, Laura Vives, Brian J. O’Roak, Peter H. Sudmant, Jay Shendure, Mark Abney, Carole Ober, and Evan E. Eichler. Estimating the human mutation rate using autozygosity in a founder population. Nature Genetics, 44:1277–1281, Sep 2012.

[15] Vagheesh M. Narasimhan, Raheleh Rahbari, Aylwyn Scally, Arthur Wuster, Dan Mason, Yali Xue, John Wright, Richard C. Trembath, Eamonn R. Maher, David A. van Heel, Adam Auton, Matthew E. Hurles, Chris Tyler-Smith, and Richard Durbin. Estimating the human mutation rate from autozygous segments reveals population differences in human mutational processes. Nature Communications, 8(1):303, 2017.

[16] Raheleh Rahbari, Arthur Wuster, Sarah J. Lindsay, Robert J. Hardwick, Ludmil B. Alexandrov, Saeed Al Turki, Anna Dominiczak, Andrew Morris, David Porteous, Blair Smith, Michael R. Stratton, UK 10K Consortium, and Matthew E. Hurles. Timing, rates and spectra of human germline mutation. Nature Genetics, 48:126–133, Dec 2015. Article.

[17] F Baudat, J Buard, C Grey, A Fledel-Alon, C Ober, M Przeworski, G Coop, and B De Massy. PRDM9 is a major determinant of meiotic recombination hotspots in humans and mice. Science, 327(5967):836–840, 2010.

[18] Augustine Kong, Gudmar Thorleifsson, Daniel F Gudbjartsson, Gisli Masson, Asgeir Sigurdsson, Aslaug Jonasdottir, G Bragi Walters, Adalbjorg Jonasdottir, Arnaldur Gylfason, Kari Th Kristinsson, et al. Fine-scale recombination rate differences between sexes, populations and individuals. Nature, 467(7319):1099–1103, 2010.

[19] Christopher L Campbell, Nicholas A Furlotte, Nick Eriksson, David Hinds, and Adam Auton. Escape from crossover interference increases with maternal age. Nature communications, 6, 2015.

[20] Amy L Williams, Giulio Genovese, Thomas Dyer, Nicolas Altemose, Katherine Truax, Goo Jun, Nick Patterson, Simon R Myers, Joanne E Curran, Ravi Duggirala, et al. Non-crossover gene conversions show strong GC bias and unexpected clustering in humans. eLife, 4:e04637, 2015.

[21] Jeffrey Staples, David J Witherspoon, Lynn B Jorde, Deborah A Nickerson, Jennifer E Below, Chad D Huff, University of Washington Center for Mendelian Genomics, et al. PADRE: Pedigree-aware distant-relationship estimation. The American Journal of Human Genetics, 99(1):154–162, 2016.

[22] Amy Ko and Rasmus Nielsen. Composite likelihood method for inferring local pedigrees. PLOS Genetics, 13(8):1–21, 08 2017.

[23] Monica D Ramstetter, Thomas D Dyer, Donna M Lehman, Joanne E Curran, Ravindranath Duggirala, John Blangero, Jason G Mezey, and Amy L Williams. Benchmarking relatedness inference methods with genome-wide data from thousands of relatives. Genetics, 207(1):75–82, 2017.

[24] Elizabeth A Thompson. Identity by descent: variation in meiosis, across genomes, and in populations. Genetics, 194(2):301–326, 2013.

[25] WG Hill and BS Weir. Variation in actual relationship as a consequence of Mendelian sampling and linkage. Genetics Research, 93(01):47–64, 2011.

[26] Brian L Browning and Sharon R Browning. Improving the accuracy and efficiency of identity-by-descent detection in population data. Genetics, 194(2):459–471, 2013.

[27] Ani Manichaikul, Josyf C Mychaleckyj, Stephen S Rich, Kathy Daly, Michèle Sale, and Wei-Min Chen. Robust relationship inference in genome-wide association studies. Bioinformatics, 26(22):2867–2873, 2010.

[28] Michael P Epstein, William L Duren, and Michael Boehnke. Improved inference of relationship for pairs of individuals. American Journal of Human Genetics, 67(5):1219–1231, 2000.

[29] Karl W Broman and James L Weber. Characterization of human crossover interference. The American Journal of Human Genetics, 66(6):1911–1926, 2000.

[30] EA Housworth and FW Stahl. Crossover interference in humans. The American Journal of Human Genetics, 73(1):188–197, 2003.

[31] International Multiple Sclerosis Genetics Consortium, Wellcome Trust Case Control Consortium 2, et al. Genetic risk and a primary role for cell-mediated immune mechanisms in multiple sclerosis. Nature, 476(7359):214–219, 2011.

[32] Kelly A Frazer, Dennis G Ballinger, David R Cox, David A Hinds, Laura L Stuve, Richard A Gibbs, John W Belmont, Andrew Boudreau, Paul Hardenbol, Suzanne M Leal, et al. A second generation human haplotype map of over 3.1 million SNPs. Nature, 449(7164):851–861, 2007.

[33] Sharon R Browning and Brian L Browning. Rapid and accurate haplotype phasing and missing-data inference for whole-genome association studies by use of localized haplotype clustering. The American Journal of Human Genetics, 81(5):1084–1097, 2007.

[34] Christopher C Chang, Carson C Chow, Laurent CAM Tellier, Shashaank Vattikuti, Shaun M Purcell, and James J Lee. Second-generation PLINK: rising to the challenge of larger and richer datasets. Gigascience, 4(1):1, 2015.

[35] Kuruvilla Joseph Abraham and Clara Diaz. Identifying large sets of unrelated individuals and unrelated markers. Source code for biology and medicine, 9(1):1, 2014.

[36] Claude Bhérer, Christopher L Campbell, and Adam Auton. Refined genetic maps reveal sexual dimorphism in human meiotic recombination at multiple scales. Nature Communications, 8, 2017.

[37] Jeffrey Staples, Dandi Qiao, Michael H Cho, Edwin K Silverman, Deborah A Nickerson, Jennifer E Below, University of Washington Center for Mendelian Genomics, et al. PRIMUS: rapid reconstruction of pedigrees from genome-wide estimates of identity by descent. American Journal of Human Genetics, 95(5):553–564, 2014.

[38] Hong Li, Gustavo Glusman, Hao Hu, et al. Relationship estimation from whole-genome sequence data. PLOS Genetics, 10(1), 2014.

[39] Alexander Gusev, Jennifer K Lowe, Markus Stoffel, Mark J Daly, David Altshuler, Jan L Breslow, Jeffrey M Friedman, and Itsik Pe’er. Whole population, genome-wide mapping of hidden relatedness. Genome Research, 19(2):318–326, 2009.

[40] Chad D Huff, David J Witherspoon, Tatum S Simonson, Jinchuan Xing, W Scott Watkins, Yuhua Zhang, Therese M Tuohy, Deborah W Neklason, Randall W Burt, Stephen L Guthery, et al. Maximum-likelihood estimation of recent shared ancestry (ERSA). Genome Research, 21(5):768–774, 2011.

[41] Braxton D Mitchell, Candace M Kammerer, John Blangero, Michael C Mahaney, David L Rainwater, Bennett Dyke, James E Hixson, Richard D Henkel, R Mark Sharp, Anthony G Comuzzie, et al. Genetic and environmental contributions to cardiovascular risk factors in Mexican Americans. Circulation, 94(9):2159–2170, 1996.

[42] Ravindranath Duggirala, John Blangero, Laura Almasy, Thomas D Dyer, Kenneth L Williams, Robin J Leach, Peter O’Connell, and Michael P Stern. Linkage of type 2 diabetes mellitus and of age at onset to a genetic location on chromosome 10q in Mexican Americans. American Journal of Human Genetics, 64(4):1127–1140, 1999.

[43] Kelly J Hunt, Donna M Lehman, Rector Arya, Sharon Fowler, Robin J Leach, Harald HH Göring, Laura Almasy, John Blangero, Tom D Dyer, Ravindranath Duggirala, et al. Genome-wide linkage analyses of type 2 diabetes in Mexican Americans. Diabetes, 54(9):2655–2662, 2005.

[44] Po-Ru Loh, Petr Danecek, Pier Francesco Palamara, Christian Fuchsberger, Yakir A Reshef, Hilary K Finucane, Sebastian Schoenherr, Lukas Forer, Shane McCarthy, Goncalo R. Abecasis, Richard Durbin, and Alkes L Price. Reference-based phasing using the haplotype reference consortium panel. Nat Genet, 48(11):1443–1448, Nov 2016.

